# Choriodecidual *Ureaplasma parvum* infection induces fetal lung inflammation prior to intra-amniotic infection in a nonhuman primate model

**DOI:** 10.64898/2026.06.06.730594

**Authors:** Sudeshna Tripathy, Nathan Crilley, Terry K. Morgan, Robert L. Schelonka, Meredith A. Kelleher

**Affiliations:** Division of Reproductive and Developmental Sciences, Oregon National Primate Research Center, Oregon Health and Science University, Beaverton OR; Division of Comparative Medicine, Oregon National Primate Research Center, Oregon Health and Science University, Beaverton OR; Department of Obstetrics and Gynecology, Oregon Health and Science University, Portland, OR; Department of Pediatrics, Division of Neonatology, Oregon Health and Science University, Portland, OR

## Abstract

Preterm birth before 28 weeks remains a leading cause of neonatal mortality and long-term morbidity. Intrauterine infection-driven chorioamnionitis is strongly associated with preterm labor, fetal inflammatory response syndrome, and neonatal lung disease. *Ureaplasma* species are among the most common organisms isolated in chorioamnionitis and are frequently detected in the placenta, amniotic fluid, and respiratory tract of preterm infants. Clinical and experimental data implicate *Ureaplasma* exposure in early lung inflammation, impaired alveolar development, and bronchopulmonary dysplasia (BPD), yet the pathogenic events preceding microbial invasion of the amniotic cavity or fetal tissues remain poorly defined.

To characterize early intrauterine and fetal lung inflammatory responses to localized choriodecidual *U. parvum* infection, we used a chronically catheterized pregnant rhesus macaque (*Macaca mulatta*) model. Time-mated animals underwent surgical placement of maternal, amniotic, and choriodecidual catheters and were inoculated with low-passage *U. parvum* serovar 1 or vehicle control at approximately 117 days gestational age. Placenta, fetal membranes, fetal plasma, and fetal lungs were assessed by qRT-PCR, multiplex cytokine assays, immunoblotting, immunohistochemistry, and trichrome staining to evaluate inflammatory signaling, inflammasome activation, prostaglandin pathways, immune cell infiltration, fibrosis, and lung maturation markers. Choriodecidual infection was confirmed in all inoculated animals. Amniotic fluid remained culture-and PCR-negative, and fetal lungs were largely free of detectable bacterial DNA. Despite the absence of intra-amniotic infection, fetal lung cytokine profiling revealed broad pro-inflammatory activation, with elevated GM-CSF, IL-1β, IL-6, IL-8, MIP-1α/β, MCP-1, VEGF and reduced IL-10 contrasting with a modest systemic response limited to elevated plasma IL-18. Fetal lungs showed increased immune cell infiltration, upregulation of NLRP3, PYCARD, and CASP1, and activation of SAPK/JNK and NF-κB signaling. Histopathology demonstrated increased alveolar macrophages and intra-alveolar neutrophils with minimal fibrosis. Surfactant gene expression was altered (increased *SFTPA*, decreased *SFTPB*), and elevated α-SMA indicated early myofibroblast activation.

Localized choriodecidual *U. parvum* infection induces fetal lung inflammation prior to detectable intra-amniotic infection, demonstrating that direct infection of the amniotic fluid or fetal lung is not required for the initiation of fetal pulmonary inflammation. These findings suggest that subclinical ascending infection may initiate fetal lung injury and increase susceptibility to postnatal respiratory morbidity associated with preterm birth.

## INTRODUCTION

Preterm birth before 28 weeks of gestation remains a leading cause of perinatal mortality and life-long morbidity worldwide. Chorioamnionitis, caused by intrauterine infection and inflammation is associated with preterm labor, premature rupture of membranes, fetal inflammatory response syndrome, cerebral palsy, and neonatal pulmonary morbidity [1–6]. Ureaplasma species are among the most commonly isolated microorganisms in chorioamnionitis and preterm labor [7–9], and have been detected in the placenta, amniotic fluid, and respiratory tract of preterm infants [10–13].

Ureaplasma respiratory tract colonization in preterm infants can occur in the context of intrauterine infection, and is strongly associated with histologic chorioamnionitis, fetal vasculitis, elevated neonatal white blood cell counts, and ruptured membranes. Colonization rates are inversely proportional to gestational age (65% at <26 weeks versus 31% at >26 weeks) [14], reflecting the compounding vulnerability of the most immature infants. Both respiratory tract colonization and the broader intrauterine inflammatory environment have been linked to chronic lung disease of prematurity called bronchopulmonary dysplasia (BPD). Among Ureaplasma species, U. parvum has been specifically associated with cardiopulmonary immaturity and respiratory distress syndrome (RDS) in neonatal respiratory samples [15], highlighting its relevance to extremely premature infant lung disease.

At the cellular level, neonatal Ureaplasma infection elicits a pronounced lower respiratory tract inflammatory response, including elevated TNF-α, IL-1β, IL-6, and IL-8, reduced anti-inflammatory IL-10, and interstitial lymphocytic infiltration [16–18]. The resulting inflammatory cascade impairs lung development, though some responses, such as increased surfactant production may transiently accelerate functional lung maturation; long-term outcomes remain unclear.

Consistent with these clinical observations, animal studies in sheep, baboons, and rhesus macaques demonstrate that prenatal Ureaplasma exposure induces inflammatory cell recruitment, profibrotic signaling, premature surfactant production, and disruption of alveolar structure [19–23], suggesting that lung injury is initiated in utero and may amplify susceptibility to subsequent postnatal insults. Despite this evidence, the early pathogenic events linking localized intrauterine infection and perinatal lung Ureaplasma infection to the severity of lung injury remain incompletely understood. Fetal and neonatal lung outcomes vary widely and are shaped by gestational age, extent and location of infection, strain-specific pathogenicity, host immune regulation, and postnatal exposures such as oxygen therapy and mechanical ventilation.

To address this gap, we employed a nonhuman primate (NHP) model of chronic choriodecidual U. parvum infection [24], in which infection is confined to the fetal membranes without amniotic fluid involvement. This model captures an intermediate stage of ascending infection to interrogate the early fetal inflammatory events that occur *in utero,* which may mediate vulnerability to subsequent lung injury and BPD. Using this model, we have previously demonstrated that chorioamnionic membranes show increased expression of inflammasome-related genes and proteins involving IL-1β and IL-18 signaling cascades in inflammatory processes and membrane weakening before the onset of preterm labor [24].

Building on these findings, the present study aimed to characterize early intrauterine and fetal lung responses to localized choriodecidual U. parvum infection in the chronically catheterized pregnant rhesus macaque (*Macaca mulatta*). We evaluated lung inflammation, inflammasome activation, and downstream kinase signaling, and assessed markers of structural remodeling including lung fibrosis (trichrome staining), α-smooth muscle actin (α-SMA) expression as a marker of alveolar septation and surfactant expression as an index of functional lung maturation.

## METHODS

### Animals

All animal procedures were approved by the Oregon Health & Science University Institutional Animal Care and Use Committee (IACUC protocol IP00000601) and conducted in accordance with the National Institutes of Health *Guide for the Care and Use of Laboratory Animals*. Animals were housed and cared for at the Oregon National Primate Research Center (ONPRC), an AAALAC International-accredited facility, under the supervision of board-certified laboratory animal veterinarians. NHPs were housed in within a climate-controlled facility, with a 12hr light/dark cycle and provided with daily environmental enrichment, including foraging puzzles, varied food treats, novel objects and visual and auditory stimulation. Veterinary staff monitored animal health and well-being daily. Animals were assigned from the ONPRC breeding colony and ovarian hormones were measured so that animals could undergo time-mated breeding. Pregnant rhesus monkeys (*Macaca mulatta*) were then adapted to a vest and mobile catheter protection system before intra-uterine surgery was performed under approved anesthetic protocols to implant amniotic, choriodecidual and maternal femoral artery and vein catheters, as previously published [21, 24–28]. Intra- and post-operative medication included analgesia (Hydromorphone and Buprenorphine), intravenous prophylactic antibiotics (Cefazolin sodium, Apotex Corp) and post-operative tocolytic medications (Terbutaline sulfate, Letco by Fagron; Atosiban, Bachem AG) as previously described. At the completion of these studies, dams were returned to the breeding colony, and fetuses were euthanized in accordance with AVMA guidelines for humane euthanasia under approved IACUC protocols.

### Experimental design

As previously published [21, 27, 29], a low-passage clinical isolate of *Ureaplasma parvum* (serovar 1) which was isolated from a case of preterm labor was provided by the Diagnostic Mycoplasma Laboratory, University of Alabama at Birmingham (UAB), USA. Pregnant animals were divided into two groups: vehicle (Control, n=4) and infected (*Ureaplasma, U.p.,* n=4). One additional animal per group was excluded from the study due to suspected preterm premature rupture of membranes (PPROM) and a secondary *Staphylococcus* infection in the amniotic fluid, respectively, as these conditions would have confounded study results. All animals underwent catheterization surgery at 108 ± 2.6 days gestational age (dGA; term = 168 days). Control animals received inoculation with sterile 2SP media or saline only and U.p. animals received *Ureaplasma parvum* inoculation via indwelling choriodecidual catheters (1 mL of 10^7^ CFU/mL in 2SP media) at 117 ± 1.6 dGA. Serial amniotic fluid samples were collected at regular intervals via indwelling amniotic catheters and sent to UAB Mycoplasma Laboratory for detection of *U. parvum* by culture and PCR, as previously published [24]. To monitor general animal health, maternal blood samples were submitted to the ONPRC Clinical Pathology Laboratory for routine chemistry panel and CBC and blood and amniotic fluid samples were submitted for general and facultative culture. Following an inoculation-to-delivery interval of 21 ± 2.4 days, animals underwent scheduled cesarean section at an average gestational age of 139 ± 1.6 dGA [24]. There were no significant differences in gestational age or inoculation-to-delivery interval between the two groups.

### Euthanasia, Necropsy, and Tissue Collection

Following scheduled cesarean section delivery, placentas were collected, weighed, cleared of blood, and trimmed of maternal decidua and fetal membrane. Placental tissue was either flash frozen in liquid nitrogen for molecular analyses or formalin fixed for histological processing.

Fetal blood was collected immediately following euthanasia into EDTA tubes, centrifuged at 4,400 rpm for 10 minutes at 4°C, and plasma was stored at -80°C. Necropsy was performed by a board-certified veterinary pathologist. The lungs were removed and the left lung inflated via tracheal cannulation using a constant hydrostatic pressure (25cm) of 10% neutral buffered formalin (NBF). Once fully distended, the bronchus was ligated to maintain inflation, and the lung was immersion-fixed in 10% NBF for ∼24hrs. Following fixation, the tissue was cut into horizontal sections and processed for paraffin embedding, with approximately six lung pieces oriented per block. The right lung was collected under sterile conditions, minced into small segments, and snap-frozen in liquid nitrogen for storage at -80°C for subsequent molecular analyses.

### Detection of Ureaplasma parvum Infection

Infection in amniotic fluid and fetal membranes was confirmed by quantitative culture and PCR for *Ureaplasma parvum* performed by the UAB Mycoplasma Lab as previously published [24]. Prior to Loop-mediated isothermal amplification (LAMP) detection of Ureaplasma DNA in fetal lung and placental samples, genomic DNA was purified from snap frozen placenta and right lung tissue stored at -80°C using a QIAamp® DNA mini kit (QIAGEN, Maryland, USA), according to manufacturer’s protocol. The isolated DNA was quantified using NanoDrop ND-1000 UV-VIS spectrophotometer (NanoDrop, Thermo Scientific, Wilmington, DE, USA) and used in reaction for *Ureaplasma parvum* detection in fetal lung samples. LAMP reactions were performed in a 25𝛍L, 2X WarmStart Multipurpose LAMP/RT-LAMP master mix (with UDG) (New England Biolabs, Ipswich, MA, USA), 10X LAMP primer mix and target DNA. The LAMP primer mix were used at the following concentrations: 1.6𝛍M (each) FIP and BIP, 0.2𝛍M (each) F3 and B3 and 0.4𝛍M (each) Loop F and Loop B. The primers were used from Fuwa et al., 2018 [30] and details are shown in S1 Table. The target gene was *ureaseB* from *Ureaplasma parvum.* The mixture was incubated at 65°C for 60 minutes and then heated at 82°C for 5 minutes for *Bst* Polymerase inactivation, in turn terminating the reaction. The presence or absence of LAMP reaction products were identified on a 1.2% agarose gel electrophoresis in 1X TAE buffer and stained with SYBR Safe stain (Invitrogen, Thermo Fisher Scientific Inc., Waltham, MA, USA). The LAMP products were run along with a GeneRuler 100 bp DNA ladder (Thermo Fisher Scientific Inc., Waltham, MA, USA). DNA extracted from a placenta positive by culture and PCR [31] for the same U. parvum serovar used in this study served as a positive control for all LAMP assays.

### Ribonucleic Acid Isolation and Quantitative Real-Time PCR

Total RNA was isolated from snap-frozen right lung tissue stored at -80°C using QIAZOL^®^ Lysis reagent (Qiagen, Maryland, USA). The fetal lung lysis was followed by ZYMO Direct-zol RNA Mini Prep Plus kit (ZYMO Research, Irvine, CA, USA), according to manufacturer’s instructions. The eluted RNA was quantified using NanoDrop ND-1000 UV-VIS spectrophotometer (NanoDrop, Thermo Scientific, Wilmington, DE, USA). RNA samples with A260/A280 values of ∼1.8 -2.0 were considered acceptable for downstream analyses.

RNA was reverse transcribed using Applied Biosystems^TM^ High-Capacity cDNA Reverse Transcription kit (Thermo Fisher Scientific), according to manufacturer’s instructions. Diluted cDNA samples equivalent to 10ng of total RNA were subjected to validation analysis on Applied Biosystems QuantStudio 3 (Thermo Fisher Scientific) employing PowerUp^TM^ SYBR Green Master mix and 2.5mM each of forward and reverse gene specific primers. Primers were designed from *Macaca mulatta* sequences in NCBI using Primer3 and BLAST, covering the exon-exon junctions. The primer details are shown in S2 Table. Expression levels of the selected genes were normalized by using *GAPDH* expression levels as internal control.

### Preparation of Tissue Lysates

Fetal lung tissue lysates were prepared in RIPA buffer (Sigma-Aldrich, St. Louis, MO, USA), supplemented with phosphatase inhibitor, PhosSTOP EASY pack (Roche, Mannheim, Germany) and Protease inhibitor cocktail EDTA free (Abcam, Cambridge, UK). Homogenized lysates were incubated on a shaker for 30 min at 4°C before centrifugation at 15,000 rpm for 15 min 4°C. The pellet was discarded, and the supernatant was recovered, aliquoted and stored at -80°C. Protein concentration was determined by colorimetric bicinchoninic acid (BCA) protein assay (Pierce, Rockford, IL, USA).

### Bead-based Multiplex Cytokine Assay

Fetal plasma and fetal lung lysate samples were submitted to the ONPRC/OHSU Endocrine Technologies Core for multiplex cytokine and chemokine quantification using Luminex technology (Millipore Sigma, Burlington, MA, USA). The following mediators were examined: GM-CSF, IL-1β, IL-1RA, IL-6, IL-8, MIP-1α, MIP-1β, MCP-1, TNF-α, VEGF, IL-18, and IL-10.

### Immunoblot analysis

Protein loading samples were resolved by SDS-PAGE on Novex™ WedgeWell™ 4–20% Tris-glycine gels (Invitrogen) and electroblotted onto nitrocellulose membranes using the iBlot 2 dry blotting system (Invitrogen, Thermo Fisher Scientific). Membranes were blocked with 5% non-fat dry milk in 1X TBST [10mM Tris pH 7.5, 150mM NaCl, 0.05% Tween-20] for 1 hour at room temperature, then incubated overnight with primary antibodies (details in S3 Table). GAPDH was used as a loading control. Membranes were incubated with HRP-conjugated secondary antibody (anti-rabbit or anti-mouse IgG, 1:5,000 in 1X TBST with 5% milk) for 1 hour at room temperature, developed with SuperSignal™ West Pico Plus chemiluminescent substrate (Thermo Scientific), and imaged on an iBright 1500 system (Invitrogen). Band densities were quantified using FIJI ImageJ 1.53k (Wayne Rasband and contributors, NIH).

### Histopathology and immunohistochemistry

The paraffin embedded blocks were cut into 5𝜇m sections and mounted on slides for immunohistochemistry (IHC) studies. The fetal lung slides were stained with hematoxylin and eosin (H&E), observed and photographed using a Carl Zeiss Axio Scope or an Olympus BX46 upright microscope with attached Olympus DP20 camera. Hematoxylin and eosin (H&E)-stained sections were examined and scored by a board-certified veterinary pathologist (NC) blinded to treatment group, using a previously established scoring system [18]. Briefly, two lung sections per animal were examined at 200x and 400x magnification for alveolar macrophages, intra-alveolar neutrophils, and lymphocytic infiltrates within alveolar walls; scores for each criterion were summed to generate an overall histology score, and the scores for the two sections were averaged to generate the score per animal. All methods followed published guidelines for experimental pathology.[32]

For immunohistochemistry, sections were deparaffinized and rehydrated through graded Histo-Clear (National Diagnostics) and ethanol washes. Antigen retrieval was performed in sodium citrate buffer (10 mM Tris-base, 1 mM EDTA, 0.05% Tween-20, pH 6.0) under high pressure. Non-specific binding was blocked with 5% normal goat serum (Thermo Fisher Scientific) and an avidin/biotin blocking kit (Vector Laboratories). Sections were incubated overnight at 4°C with primary antibodies targeting CD68 (macrophages), CD45 (leukocytes), PTGS2/COX-2 (prostaglandin synthase), and α-SMA (smooth muscle actin) in fetal lung; and CD68 (M1 macrophages) and CD163 (M2 macrophages) in fetal membrane (antibody details in S3 Table). Secondary antibody incubation was followed by ABC reagent (VECTASTAIN Elite ABC HRP Kit) and developed with Vector NovaRed Substrate Kit (Vector Laboratories). Slides were scanned on an Olympus VS120 slide scanner at 20x (0.75 NA) and 40x (0.95 NA) objectives. Fetal membrane sections were scored by a placental pathologist (TKM) blinded to treatment group; M1 (CD68+) and M2 (CD163+) macrophages were counted per high-power field (400x), with the mean of four fields per section reported. Image analysis was performed using FIJI ImageJ 1.53k with background subtraction (rolling ball radius 50 pixels), color deconvolution, and area fraction quantification normalized to nuclear staining coverage; four sub-areas per section were averaged per animal.

### Trichrome Staining

Collagen deposition in fetal lung sections was assessed using Masson’s trichrome stain (IMEB Inc., CA, USA) according to the manufacturer’s instructions. Briefly, deparaffinized and rehydrated sections were pretreated with Bouin’s solution, stained sequentially with hematoxylin, Biebrich scarlet-acid fuchsin, and aniline blue, then dehydrated and mounted with DPX. Quantification of lung fibrosis was performed on scanned trichrome-stained slides using an automated image analysis model in Orbit version 3.64 [33, 34] by a board-certified veterinary pathologist (NC).

### Statistical Analyses

Data are expressed as mean ± SEM unless otherwise indicated. Normal distribution was assessed using the Shapiro-Wilk test. Between-group comparisons were performed using Student’s t-test for normally distributed data or the Mann-Whitney U-test for non-parametric data. Given the small sample size inherent to NHP studies, no correction for multiple comparisons was applied; p-values should be interpreted accordingly. A p-value of <0.05 was considered statistically significant. All analyses were performed in GraphPad Prism, version 11.01, GraphPad Software, San Diego, CA

### AI Assistance

Artificial intelligence tools (Claude, Anthropic; Microsoft Copilot) were used to assist with manuscript drafting and editing. All content was critically reviewed, revised, and approved by the authors, who take full responsibility for the accuracy and integrity of the work.

## RESULTS

### Confirmation of localized choriodecidual Ureaplasma parvum infection

As previously published, all animals in the choriodecidual infection group (*U.p.)* were confirmed to have *U. parvum* bacteria at the fetal membrane inoculation site but no detectable *Ureaplasma* in the amniotic fluid by culture and PCR assays (Table 1; [24]). To determine the extent of microbial spread beyond the membranes, placental and fetal lung DNA were analyzed using *ureaseB* LAMP. Among the four infected animals in the *U.p.* group, two showed *U. parvum*–positive placental samples (S1A Fig), and one of these also demonstrated *ureaseB*-positive fetal lung tissue (S1B Fig). As expected, all control animals were negative for *Ureaplasma* across all samples.

**Table 1.**
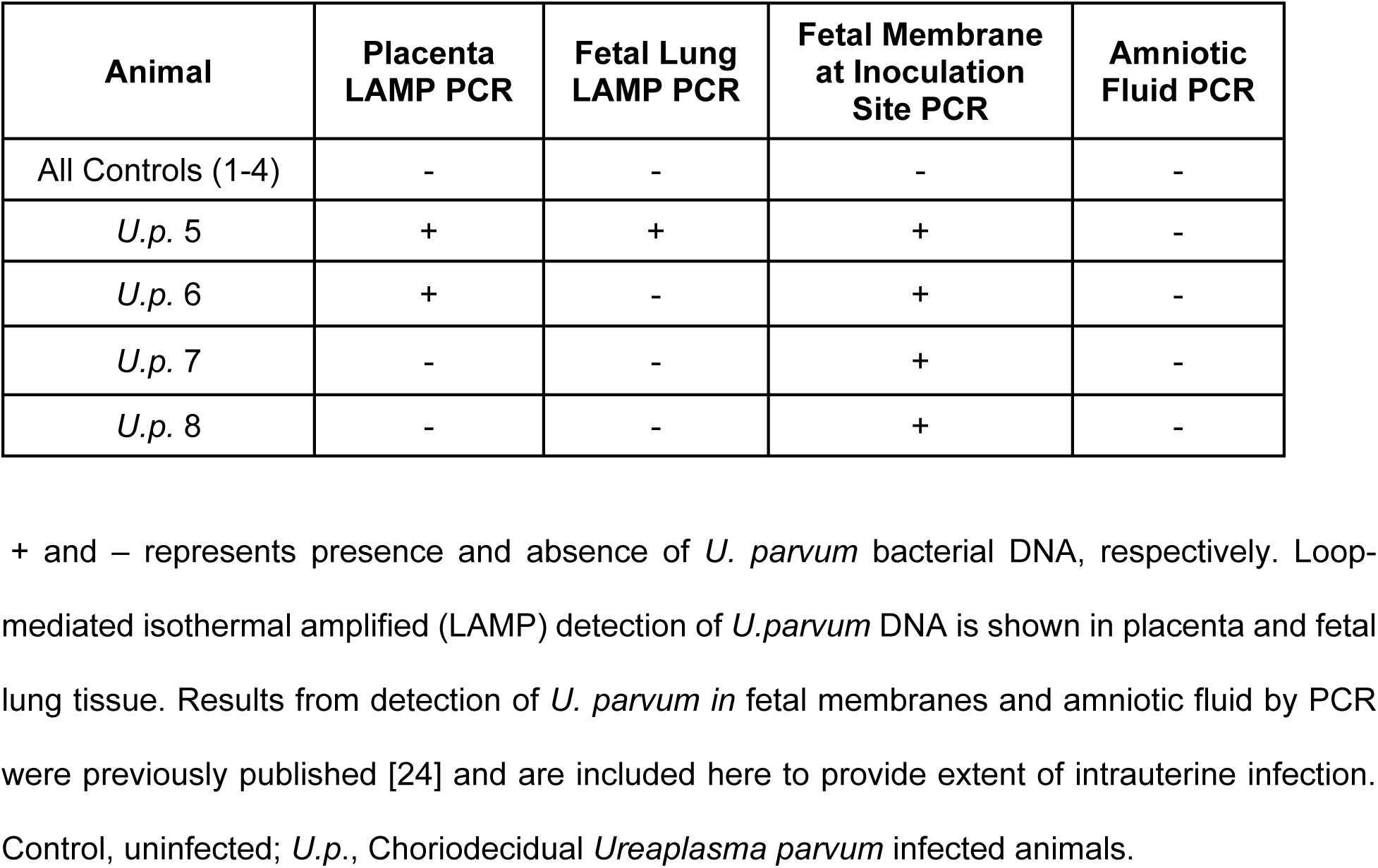
Summary of *Ureaplasma parvum* detection in amniotic fluid, fetal membranes, placenta and fetal lung.

### Choriodecidual Ureaplasma parvum infection induces inflammatory activation in placenta and fetal membranes

Placental expression of inflammatory mediators (*TNF-α, IL-6, IL-8, IL-1α, IL-1β, IL-1R1, IL-1RA, MCP-1, VEGF, IL-18*) was assessed by RT-qPCR (Fig 1A). *TNF-α, IL-1β,* and *MCP-1* mRNA were significantly elevated in placentas from animals in the *U.p.* group (p = 0.01). No significant differences in placental cytokine protein abundance were detected by multiplex analysis (S2 Fig). Immunohistochemistry of fetal membranes revealed increased pro-inflammatory M1 macrophages (CD68+) and decreased M2 macrophages (CD163+) in infected animals (Fig 1B). M1/M2 clustering indicated a predominantly moderate inflammatory phenotype in the *U.p.* group, compared with low inflammation in controls (Fig 1C).

**Fig 1.**
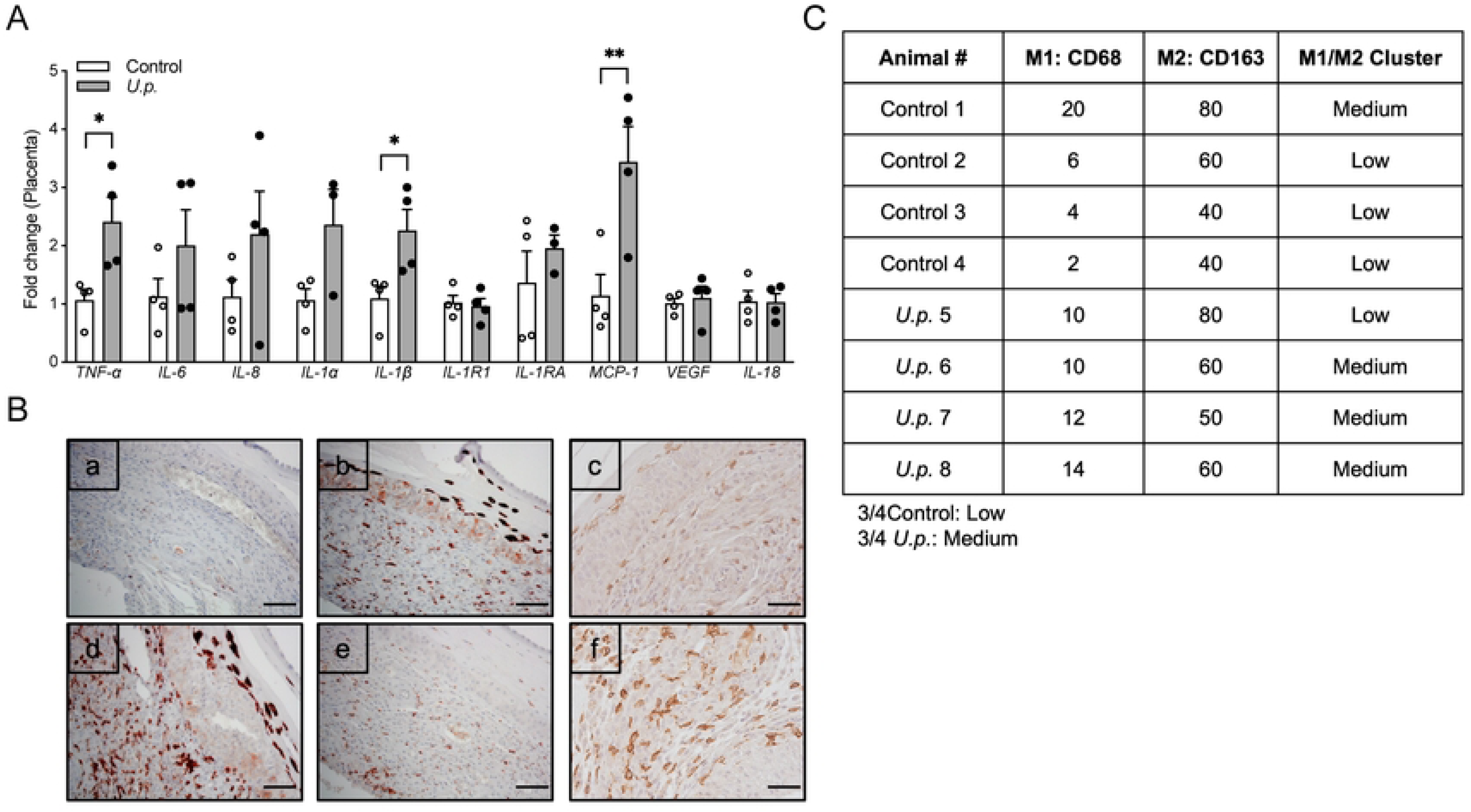
*Ureaplasma parvum* infection induced changes in inflammatory mediators in placenta and fetal membrane. (A) Placental gene expression of *TNF𝛂*, *IL-1β* and *MCP-1* was increased in animals exposed to choriodecidual *U.p.* compared to control. mRNA expression were normalized to the *GAPDH* housekeeping gene and represented as mean±SEM fold change. (B) Representative immunohistochemical staining for control (a, b, c) and *U.p.* (d, e, f) animals. M1 macrophages, CD68+ (a, d); M2, CD163+ (b, e) and leukocytes, CD45+ (c, f) are shown. (C) The number of M1(pro-inflammatory) and M2 (anti-inflammatory) macrophages (B; a, d, b, e) were scored per high power field (hpf). The mean of four separate hpf/section were reported for each case as the number of CD68+ macrophages/hpf and the number of CD163+ macrophages/hpf. The M1/M2 ratio was calculated per animal and categorized as Low (<0.15) or Medium (0.15–0.25). Control animals were predominantly Low (3/4), while *U. parvum*-inoculated animals were predominantly Medium (3/4), consistent with a shift toward greater M1 representation following choriodecidual infection. Control, uninfected; *U.p.*, Choriodecidual *Ureaplasma parvum* infected animals; hpf, high power field. Scale bar = 20𝜇m.

### Choriodecidual infection produces selective systemic inflammation in the fetus

Fetal plasma cytokines (GM-CSF, IL-1β, IL-1RA, IL-6, IL-8, MIP-1α, MIP-1β, MCP-1, TNF-α, VEGF, IL-18, IL-10) were measured by multiplex assay (Fig 2A). IL-18 was significantly elevated in *U.p.* fetuses (p = 0.02). Other pro-inflammatory mediators commonly associated with fetal inflammatory response syndrome, including TNF-α, IL-6, and IL-8, did not differ significantly between the control and *U.p.* groups.

**Fig 2.**
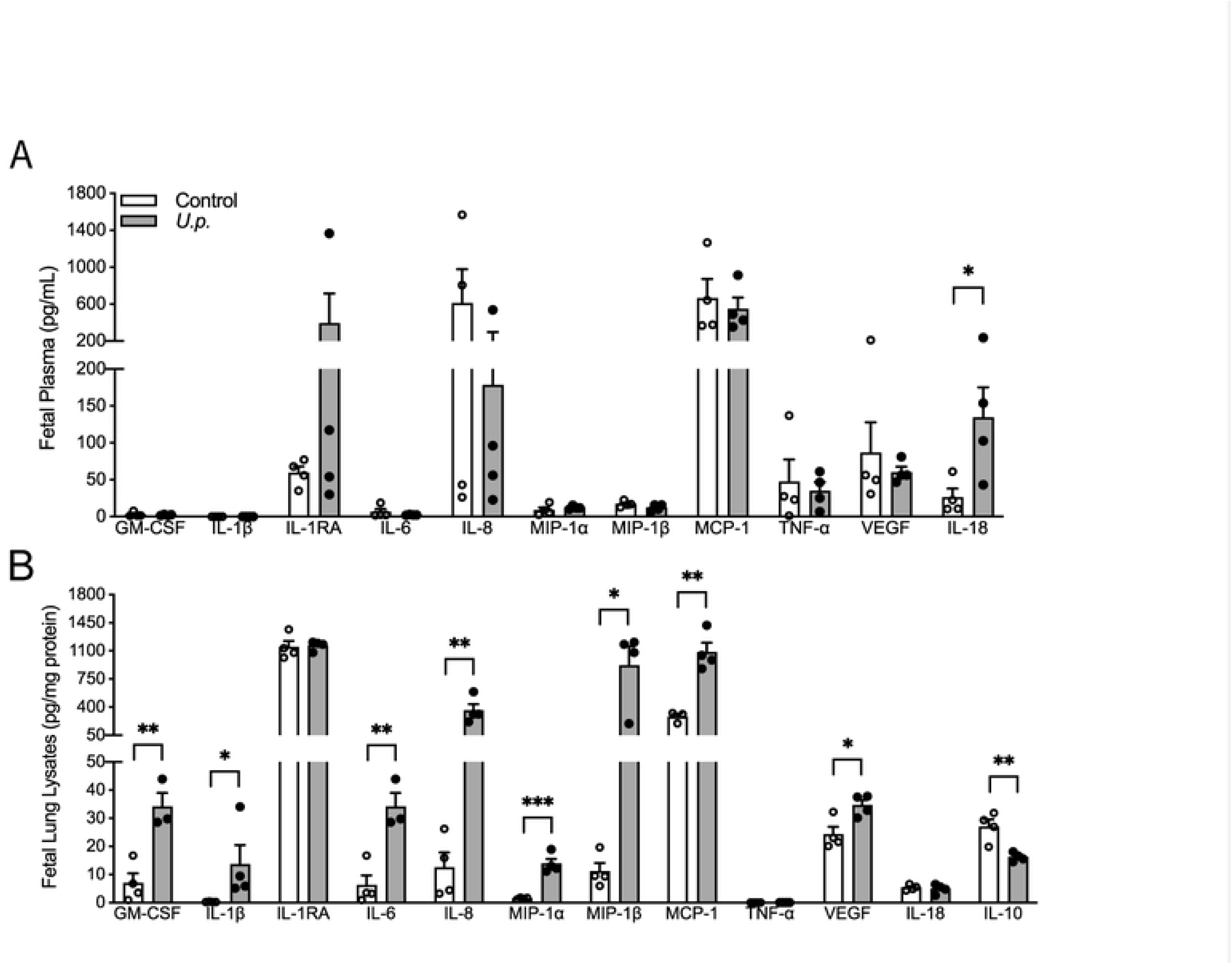
Fetal plasma and lung inflammatory mediator profiles following choriodecidual *U. parvum* infection. (A) Fetal plasma IL-18 was significantly increased in the animals exposed to choriodecidual *U.p.* compared to control. (B) In the fetal lungs, GM-CSF, IL-1β, IL-6, IL-8, MIP-1α, MIP-1β, the MCP-1 and VEGF were significantly increased, and the anti-inflammatory cytokine IL-10 was significantly reduced in the *U.p.* group compared to control. Each bar represents the mean±SEM. Asterisks indicate statistically significant difference from the corresponding control (Student’s t-test/ Mann-Whitney U-test, *P<0.05, **P<0.01 and ***P<0.005, n=3-4 animals/group). Control, uninfected; *U.p.*, Choriodecidual *Ureaplasma parvum* infected animals.

### Choriodecidual Ureaplasma parvum infection elicits robust inflammatory responses in the fetal lung

In contrast to the modest systemic response, fetal lung cytokine profiles showed widespread inflammation. GM-CSF, IL-1β, IL-6, IL-8, MIP-1α, MIP-1β, MCP-1, and VEGF protein concentrations were significantly increased, whereas IL-10 was significantly decreased in infected lungs (P < 0.05; Fig 2B). Corresponding changes in *TNF-α, IL-6, MCP-1, VEGF*, and *IL-10* mRNA were observed (S3 Fig). The fetus with ureaseB-positive lung tissue did not exhibit cytokine patterns that differed from *ureaseB*-negative infected animals.

### Fetal lung histopathology reveals inflammation with minimal fibrosis

H&E evaluation demonstrated increased alveolar neutrophils and macrophages and enhanced lymphocytic infiltration in the lungs of fetuses from the *U.p.* group (Fig 3A–B). Although the composite histopathology score trended higher in the *U.p.* group compared to control, the difference did not reach statistical significance (Fig 3C). Masson trichrome staining showed minimal collagen deposition in fetal lungs from *U.p.* animals and no substantial fibrosis compared to controls (Fig 3Ac–f).

**Fig 3.**
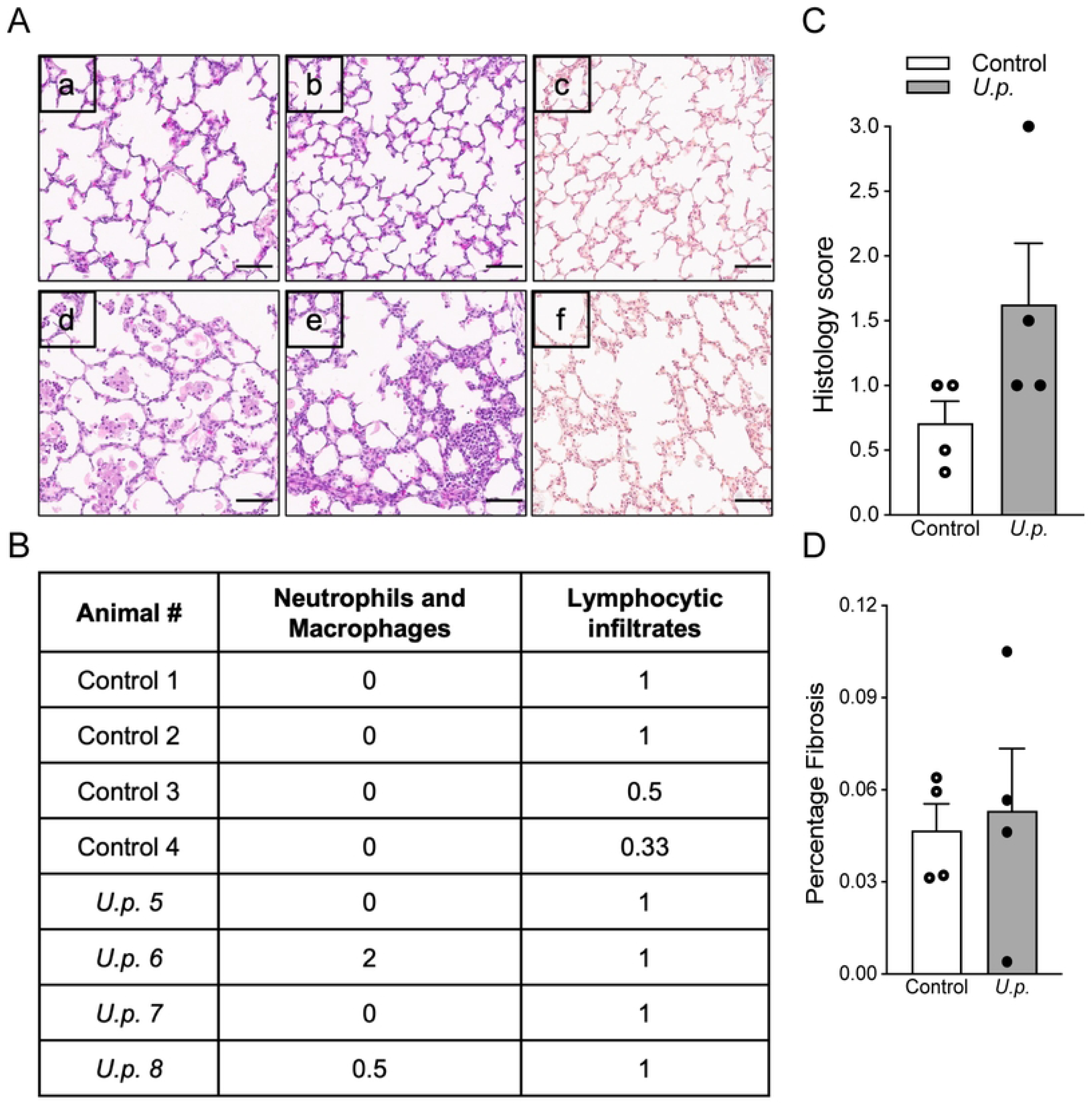
*Ureaplasma parvum* infection induced significant inflammation and minimal fibrosis in fetal lung. (A) Representative photomicrographs of lung pathology in control (a, b, c) and *U.p.* (d, e, f) animals showing H&E (a, b, d, e) and trichrome (c, f) staining. *U.p.* animal alveolar spaces contain neutrophils, macrophages and inflammatory debris (d) while alveolar walls contain moderate lymphocytic infiltrates (e) compared to control (a and b, respectively). Trichrome staining reveals minimal fibrotic changes in U.p. animals (f) compared to controls (c). (B) Table shows the numerical score for Neutrophils/Macrophages and Lymphocytic Infiltrates determined by the average score between 2-3 regions/animal. (C) Graph represents overall histology score (sum of neutrophil/macrophage and lymphocytic infiltrate scores). (D) Fibrosis quantified as the percentage of trichrome-positive area relative to total tissue area per animal. Each bar represents the mean± SEM, Student’s t-test/ Mann-Whitney U-test, n=4 animals/group. Scale bar, 75 μm. Control, uninfected; *U.p.*, Choriodecidual *Ureaplasma parvum* infected animals.

### Increased immune cell infiltration and prostaglandin pathway activation in the fetal lung

By immunohistochemistry, the staining of CD45+ leukocytes and CD68+ macrophages were significantly increased in infected fetal lungs (P<0.05; Fig 4A–C). Both PTGS2 mRNA and protein expression was significantly elevated in lungs from fetuses in the *U.p.* group when compared to controls (P<0.05; Fig 4A–C). Expression of the prostaglandin-degrading enzyme HPGD was unchanged (Fig 4A).

**Fig 4.**
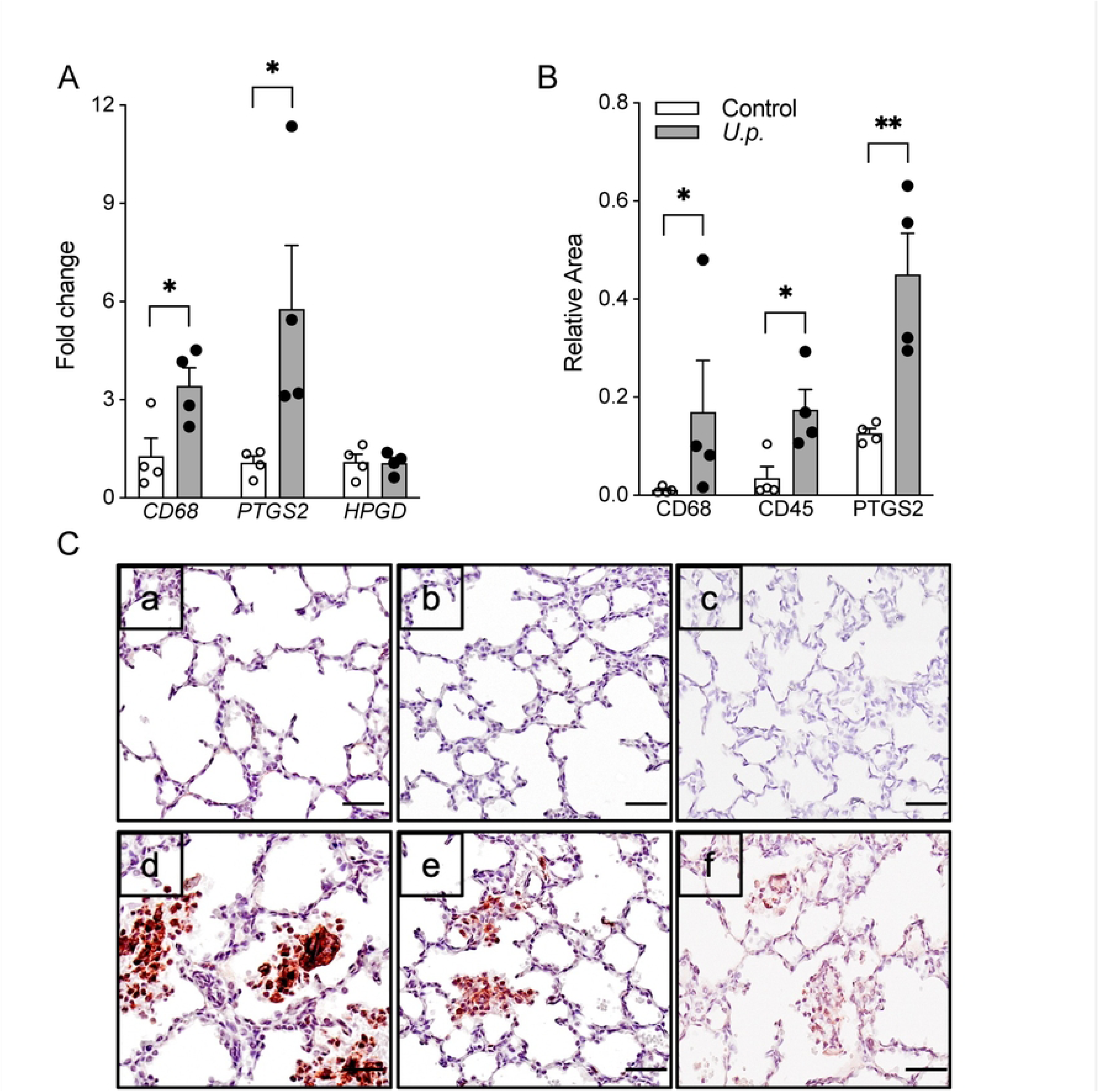
Increased inflammatory markers and prostaglandin synthesis in fetal lung. (A) mRNA expression of *CD68* and *PTGS2* was significantly increased in the fetal lungs of *U.p.* animals compared to control when normalized to *GAPDH* housekeeping gene. (B) Quantitative immunohistochemical analyses of CD68, CD45 and PTGS2 in fetal lung shows significantly increased staining in the fetal lungs of *U.p.* animals compared to controls. Staining for each protein was calculated relative to nuclear staining area. (C) Representative immunohistochemical staining are shown for control (a, b, c) and *U.p.* (d, e, f) animals for CD68 (a, d), CD45 (b, e) and PTGS2 (c, f) (Scale bar, 20 μm). Each bar represents the mean±SEM. Asterisks indicate significant difference from the corresponding control (Student’s t-test/ Mann-Whitney U-test, *P<0.05, **P<0.01, n=4 animals/group). Control, uninfected; *U.p.*, Choriodecidual *Ureaplasma parvum* infected animals.

### Upregulation of IL-18 and inflammasome components in fetal lung

*IL-18* mRNA was significantly increased in lungs from the *U.p.* group (P < 0.01; Fig 5A). Western blotting detected both pro- and mature IL-18 but did not detect any significant differences in protein expression between the two groups (Fig 5B–C). Fetal lung IL-18R1 protein expression showed a trend toward increased abundance in the *U.p.* group (p = 0.06; Fig 5D). *CASP1* mRNA was significantly elevated in infected lungs (P<0.05; Fig 5A), although CASP1 protein levels were unchanged. CASP4, IL-18BP, and IL-18RAP gene or protein expression did not differ between groups (S4 Fig).

**Fig 5.**
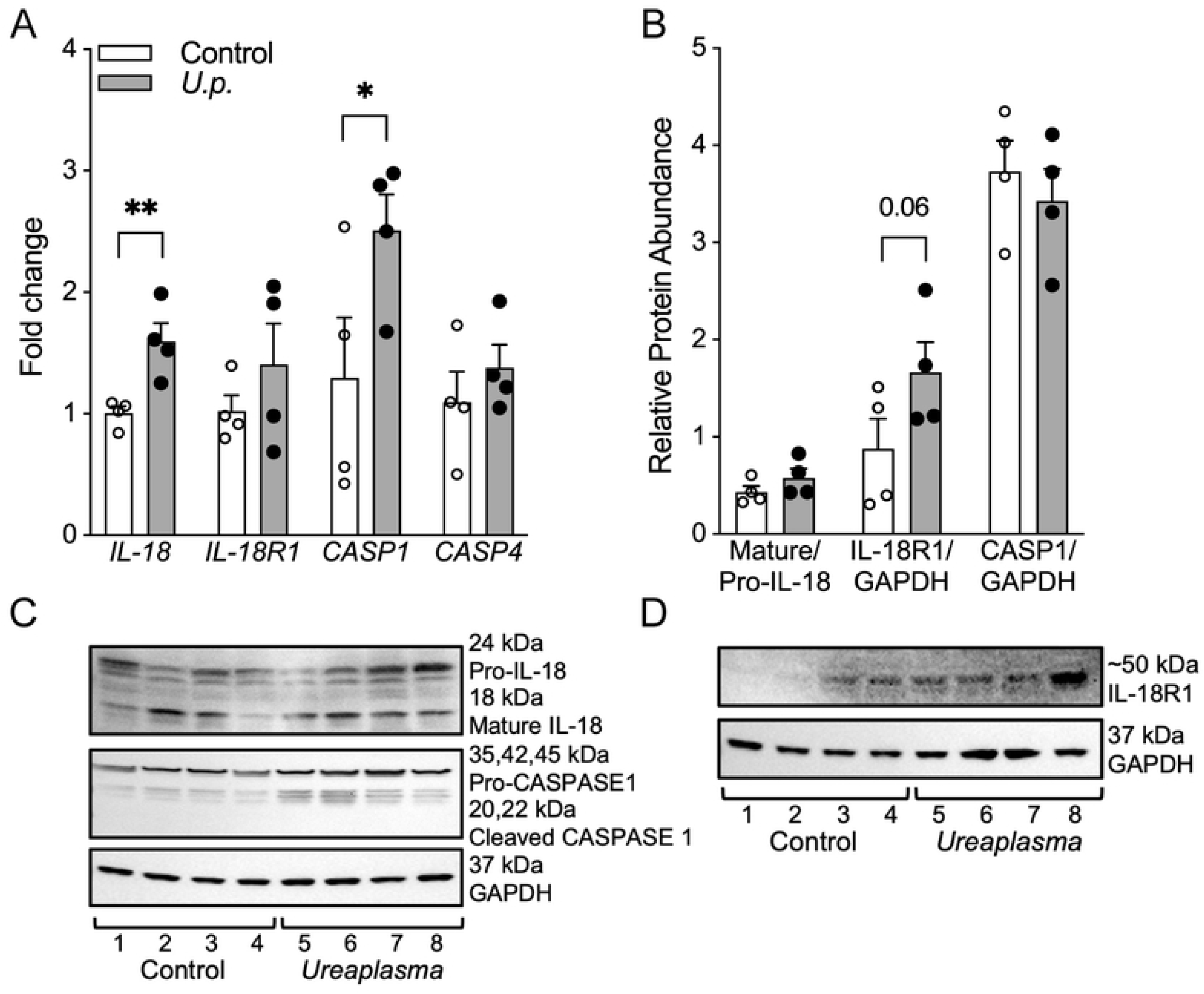
IL-18, its receptor and caspase cleaving enzyme changes in fetal lung. (A) mRNA expression of *IL-18* and C*ASP1* significantly increased in the fetal lung of *U.p.* animals compared to control when normalized to *GAPDH* housekeeping gene. (B) Densitometric quantification and (C and D) representative western blot of pro-IL-18, mature IL-18, pro-CASP1, cleaved CASP1 and IL-18R1 in the fetal lung. Target protein levels were normalized to GAPDH as a loading control. Each bar represents the mean± SEM. The asterisks indicate significant difference from the corresponding control (Student’s t-test/ Mann-Whitney U-test, *P<0.05 and **P<0.01, n=4 animals/group). Control, uninfected; *U.p.*, Choriodecidual *Ureaplasma parvum* infected animals.

Gene and protein expression of inflammasome components were significantly increased in the fetal lungs from animals in the *U.p.* group, including NLRP3 and the adaptor PYCARD/ASC-TMS1 (P < 0.05; Fig 6A–D). *NOD2* and *AIM2* mRNA were also elevated but corresponding protein levels for these were unchanged. NLRC4 expression showed no differences across groups.

**Fig 6.**
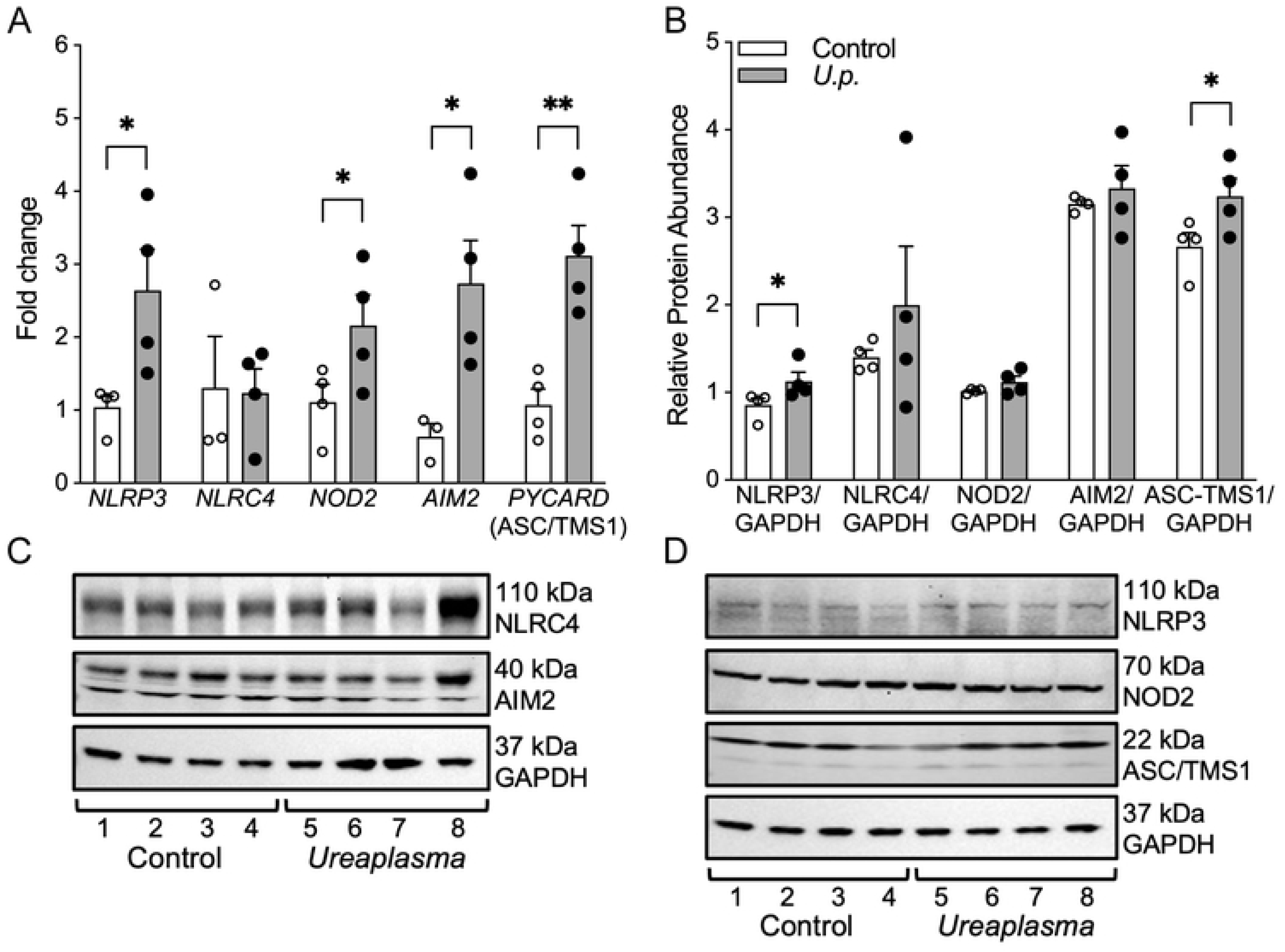
Inflammasome upregulation in fetal lung. (A) mRNA expression of inflammasome sensors *NLRP3*, *NOD2*, *AIM2* and adaptor *PYCARD* are significantly upregulated in the fetal lung of *U.p.* animals compared to control when normalized to the *GAPDH* housekeeping gene. (B) Densitometric quantification and (C and D) representative western blots of NLRP3, NLRC4, NOD2, AIM2 and ASC-TMS1 in the fetal lung of *U.p.* animals compared to control, showed a significant increase in NLRP3 and ASC-TMS1 protein expression. Target protein levels were normalized to GAPDH as a loading control. Each bar represents the mean± SEM. The asterisks indicate significant difference from the corresponding control (Student’s t-test/ Mann-Whitney U-test, *P<0.05 and **P<0.01, n=4 animals/group). Control, uninfected; *U.p.*, Choriodecidual *Ureaplasma parvum* infected animals.

### Activation of downstream IL-18 signaling kinases

Downstream signaling components of the IL-18 pathway related to NF-κB signaling were examined. *RELA* (NF-κB p65) mRNA was significantly upregulated, while *TAB2* was significantly decreased in infected lungs (P < 0.05; Fig 7A). SAPK/JNK protein activation was significantly increased (p = 0.02; Fig 7B–C). MYD88, IRAK1, ERK1/2, and p38 MAPK did not show differences between groups (S5 Fig).

**Fig 7.**
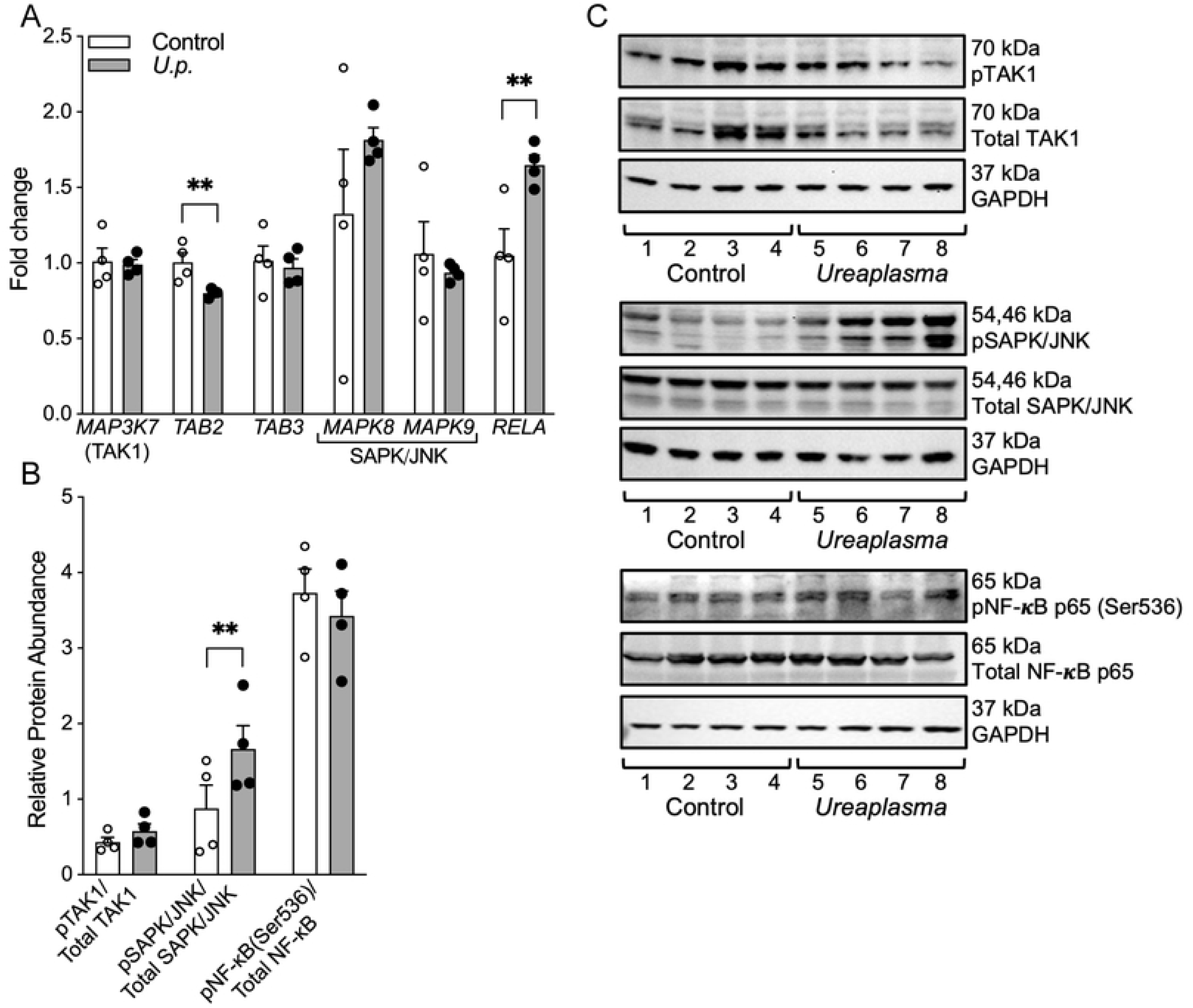
NF-κB and multiple kinase pathway changes in fetal lung. (A) mRNA expression of *RELA (*NF-κB p65 subunit) increased *and TAB2* decreased in the *U.p.* animals compared to controls when normalized to the *GAPDH* housekeeping gene. (B) Densitometric quantification and (C) representative western blot showed a significant increase in active pSAPK/JNK in the *U.p.* animals compared to controls. Target protein levels were normalized to GAPDH as a loading control. The results for each target protein are shown as the active phosphorylated form relative to total protein level. Each bar represents the mean±SEM. The asterisks indicate significant difference from the corresponding control (Student’s t-test/ Mann-Whitney U-test, **P<0.01, n=4 animals/group). Control, uninfected; *U.p.*, Choriodecidual *Ureaplasma parvum* infected animals.

### Choriodecidual infection alters surfactant expression and induces myofibroblast activation in fetal lung

Surfactant gene expression was altered in infected lungs, with a significant increase in (P<0.01) and a significant decrease in *SFTPB* (P<0.01), while *SFTPC* and *SFTPD* were unchanged (Fig 8A). Despite reduced *SFTPB* mRNA, immunohistochemistry showed increased protein localization for both surfactant A and B in infected lungs (Fig 8B–C).

**Fig 8.**
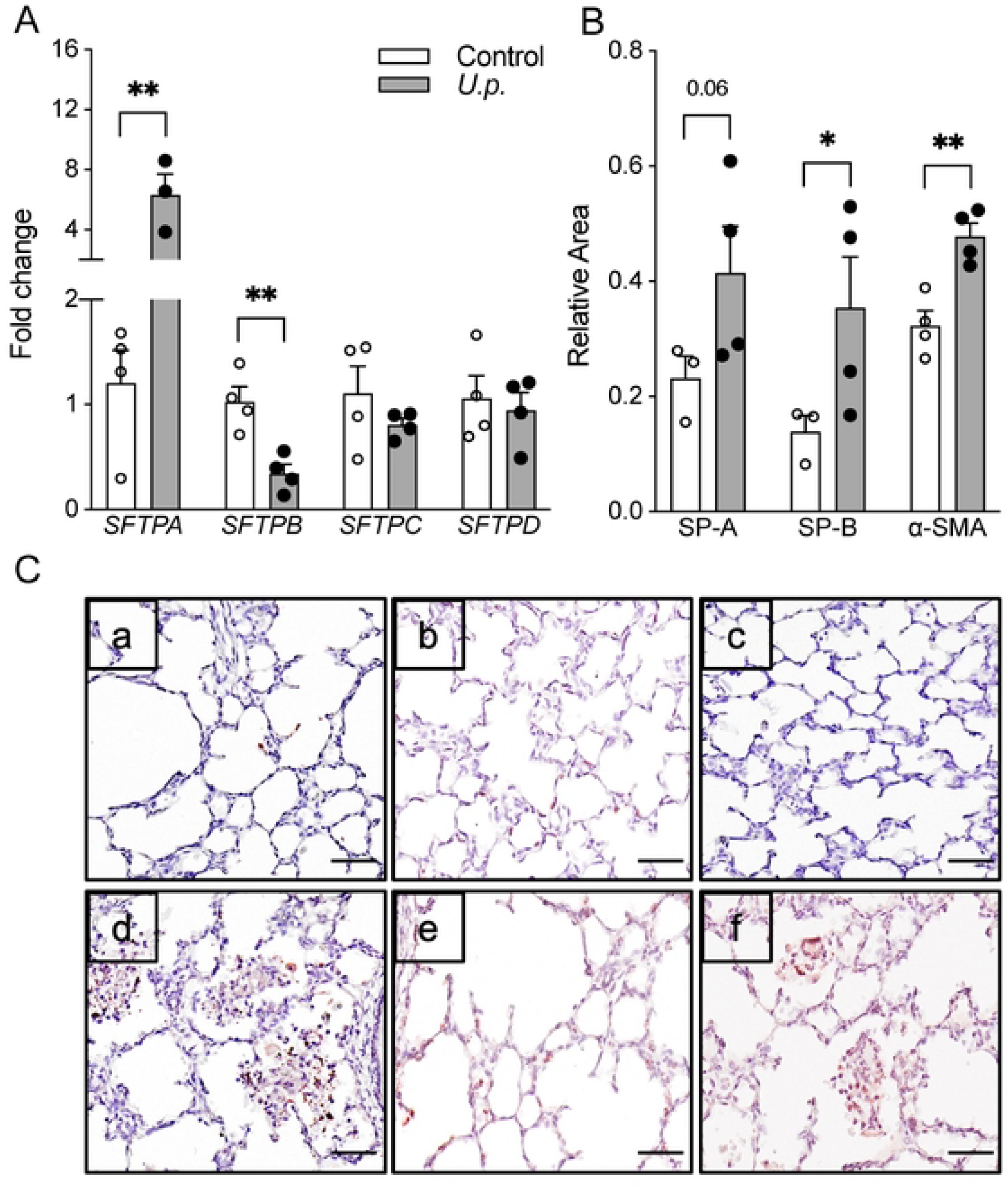
Surfactant and Alpha Smooth Muscle Actin (α-SMA) distribution in fetal lung. (A) mRNA expression of surfactant genes *SFTPA* and *SFTPB* were significantly upregulated and downregulated, respectively in the *U.p.* animals compared to control when normalized to *GAPDH* housekeeping gene. (B) Quantitative immunohistochemical analyses of SP-A, SP-B and alveoli α-SMA shows increased staining in the fetal lung for all protein in *U.p.* animals compared to control. Staining for each protein was calculated relative to nuclear staining area. (C) Representative immunohistochemical staining are shown for control (a, b, c) and *U.p.* (d, e, f) animals for SP-A (a, d), SP-B (b, e) and α-SMA protein expression in alveoli (c, f). Each bar represents the mean± SEM. The asterisks indicate significant difference from the corresponding control (Student’s t-test/ Mann-Whitney U-test, *P<0.05, **P<0.01, n=3-4 animals/group). Scale bar, 20 μm. Control, uninfected; *U.p.*, Choriodecidual *Ureaplasma parvum* infected animals.

α-Smooth muscle actin (α-SMA), a marker of myofibroblasts involved in alveolar septation and injury, was significantly increased in alveolar regions of infected lungs (P<0.05; Fig 8B, 8Cf). α-SMA levels in pulmonary arteries and bronchial airways were unchanged (S6 Fig).

## DISCUSSION

In this study, we employed a chronically catheterized rhesus macaque model to characterize early fetal lung responses to localized choriodecidual *Ureaplasma parvum* infection in the absence of intra-amniotic infection or preterm labor [24]. This model allowed us to examine inflammatory signaling and structural changes in the fetal lung during an early stage of ascending infection, before microbial invasion of the amniotic cavity or widespread fetal infection. Our findings demonstrate that even when *U. parvum* infection is confined to the choriodecidual space, the fetus mounts a robust lung inflammatory response that is disproportionate to the modest systemic response detected in fetal plasma, and which is not explained by direct amniotic fluid exposure or the extent of fetal lung infection by *Ureaplasma*. These data suggest that fetal lung injury cascade may be initiated before clinical signs of chorioamnionitis or intrauterine infection are detectable, highlighting the gap between current diagnostic capacity and the therapeutic window to improve postnatal respiratory outcomes.

Systemic cytokine elevations were minimal, limited to increased IL-18 in fetal plasma (Fig 2A). In contrast, cytokine profiling of the fetal lung revealed broad pro-inflammatory activation, including elevated GM-CSF, IL-1β, IL-6, IL-8, MIP-1α/β, MCP-1, and VEGF, alongside reduced IL-10 (Fig 2B). This profile aligns with cytokine patterns identified with bronchopulmonary dysplasia (BPD), encompassing inflammatory cell recruitment, myeloid activation, and vascular remodeling, coupled with downregulation of IL-10-mediated anti-inflammatory responses [14, 35]. The modest systemic response may partly reflect the approximately 3 week interval between inoculation and sample collection, raising the possibility that broader systemic inflammation may have been present earlier in the infection course but had subsided by the time of sampling. The isolated elevation of IL-18 is consistent with this interpretation, as inflammasome-mediated IL-18 maturation can sustain production beyond the early-phase cytokine response, potentially remaining detectable after other systemic signals have resolved [36–39]. Notably, elevated IL-18 has previously been associated with BPD development in preterm infants [40], suggesting that even this limited systemic signal may carry prognostic relevance for adverse pulmonary outcomes.

A notable finding was the coordinated upregulation of inflammasome-related genes and proteins, including NLRP3 and PYCARD/ASC-TMS1 (Fig 6), alongside elevated IL-18 mRNA (Fig 5) [24, 41–43]. NF-κB pathway activation, a key upstream regulator of NLRP3 transcription and IL-18 processing, likely contributes to this coordinated response and may represent the mechanistic link between choriodecidual infection and fetal lung inflammasome priming [44–46]. These results extend previous observations of inflammasome activation in the chorioamnionic membranes in this model following choriodecidual Ureaplasma inoculation, indicating that the fetal lung is similarly primed toward inflammasome-mediated inflammatory signaling [24]. However, significant mRNA upregulation of IL-18, CASP1, NOD2, and AIM2 was not accompanied by corresponding increases in total protein abundance for these genes. This may reflect post-transcriptional regulation, translational repression, or rapid protein turnover in the context of active inflammasome assembly [41, 47] and does not preclude functional activation of these pathways. Consistent with this interpretation, inflammasome-mediated IL-18 cleavage and secretion into the extracellular compartment may account for the absence of intracellular protein accumulation despite elevated mRNA, a pattern that warrants investigation in future studies measuring secreted IL-18 in bronchoalveolar lavage or tissue supernatants. SAPK/JNK protein activation was also significantly increased in infected fetal lungs (Fig 7). JNK signaling has been implicated in apoptosis, cytokine production, and impaired alveolar development in the context of neonatal lung injury [48, 49], and its activation here may contribute to the structural and functional alterations observed.

Although fetal lung tissue rarely contained detectable bacterial DNA, this does not preclude biologically meaningful fetal exposure via other routes. Soluble inflammatory mediators, prostaglandins, extracellular vesicles, and DAMPs produced in the decidua and membranes can traffic through the fetal circulation [50], potentially exposing the fetus to sustained pro-inflammatory signals without direct microbial invasion, and likely contribute to the strong pulmonary response observed in the present study. Supporting this, immunohistochemistry of the fetal membranes revealed a shift towards M1 macrophage (CD68+) predominance with reduced anti-inflammatory M2 (CD163+) macrophages in infected animals (Fig 1), consistent with an activated pro-inflammatory phenotype at the maternal-fetal interface, suggesting that indirect maternal-fetal signaling, rather than direct amniotic fluid exposure or fetal infection, is sufficient to drive pulmonary inflammation in the developing fetus. Unlike prior NHP studies of choriodecidual infection in which significant amniotic fluid cytokine elevation was documented [51], the pronounced fetal lung inflammatory response observed here occurred in the absence of detectable intra-amniotic infection or amniotic fluid inflammation, indicating that aspiration of inflammatory mediators is not required to initiate fetal lung inflammation. The one animal with ureaseB-positive fetal lung tissue did not exhibit a distinct cytokine profile compared to infected animals without detectable fetal lung Ureaplasma, further supporting the interpretation that the inflammatory response is driven primarily by indirect signaling rather than direct bacterial colonization of the lung parenchyma.

Despite this inflammatory environment, histopathologic evaluation revealed minimal fibrosis, consistent with the relatively short interval between inoculation and delivery and the absence of postnatal exposures, such as supplemental oxygen or mechanical ventilation, that contribute to lung injury [52–54]. Significant increases in alveolar macrophages, intra-alveolar neutrophils, and leukocytes were nonetheless observed, indicating active acute immune cellular recruitment and infiltration. Prior NHP studies using the same *Ureaplasma* clinical isolate delivered via direct intra-amniotic inoculation produced peribronchiolar lymphocytic aggregates [27], a finding absent in the current model. This difference likely reflects direct fetal lung exposure via aspiration of infected amniotic fluid in the intra-amniotic infection model, which would be expected to elicit a more intense local immune response than localized choriodecidual infection without microbial invasion of the amniotic cavity. Upregulation of PTGS2 (Fig 4) further suggests activation of prostaglandin-mediated pathways that may contribute to inflammatory signaling, tissue remodeling, or altered surfactant production [55]. Overall, these data demonstrate that significant pulmonary inflammatory responses can be established even in the absence of extensive direct fetal lung infection.

Alterations in surfactant expression, with increased SFTPA and decreased SFTPB mRNA but increased protein staining for both surfactant A and B (Fig 8), are consistent with prior evidence that intrauterine inflammation can accelerate certain aspects of functional lung maturation while simultaneously contributing to cellular stress and injury [56]. Increased α-SMA in alveolar regions (Fig 8) indicates early myofibroblast activation, a hallmark of disrupted septation and alveolarization, which are early features of BPD pathogenesis [23, 57]. The timing of exposure likely influences the nature of injury, as the fetal lung at 105–145 dGA in rhesus macaques corresponds to the late canalicular-early saccular stages, when epithelial differentiation, vascular remodeling, and surfactant expression are undergoing rapid remodeling [58]. Inflammatory activation during this developmental window may therefore have disproportionate long-term consequences compared to later gestation. Importantly, these findings demonstrate that even mild intrauterine inflammation, in the absence of direct fetal infection, is sufficient to alter pathways involved in fetal lung development and may increase vulnerability to postnatal respiratory morbidity.

Limitations of this study include the small number of animals that can be ethically and practically studied, intrinsic to nonhuman primate experiments. Despite this, NHP models offer exceptional fidelity to human reproductive and developmental biology. Rhesus macaque placentation, immune maturation, and fetal lung development closely parallel those of humans, including the timing of alveolarization, surfactant biology, and responses to in utero exposures, rendering primate models particularly valuable for reproductive and perinatal research where the complexity of maternal-fetal interactions cannot be replicated *in vitro* or *in silico* [59, 60]. The duration of infection may also be insufficient for the development of later-stage injury such as fibrosis or alveolar simplification, and the model does not incorporate postnatal exposures central to BPD pathogenesis [61, 62]. Nonetheless, isolating early intrauterine inflammatory effects in the absence of postnatal confounders or widespread fetal infection provides a unique opportunity to examine fetal lung priming during a clinically relevant developmental window that cannot be studied in human pregnancy.

Consistent with previously published findings from this model demonstrating robust inflammasome activation and pro-inflammatory cytokine upregulation in the chorioamnionic membranes prior to preterm labor [24], the current data suggest that choriodecidual *Ureaplasma* infection establishes a tissue-specific gradient of inflammation across the maternal-fetal interface. Inflammasome-mediated membrane weakening and the concurrent fetal lung inflammatory response described here may therefore represent parallel pathways through which subclinical ascending infection contributes to adverse pregnancy outcomes and inflammatory-mediated altered fetal pulmonary development, even before clinical signs of chorioamnionitis become evident.

In summary, localized choriodecidual *U. parvum* infection elicits a pronounced inflammatory response in the fetal lung despite the absence of intra-amniotic infection, fetal dissemination, or broad systemic fetal inflammatory response. These findings identify early inflammatory, inflammasome, and structural alterations that may prime the fetus for later respiratory morbidity. Inflammation is a potentially modifiable risk factor for lung injury in premature infants [63] and a target of emerging anti-inflammatory treatments for preterm labor, suggesting that interventions at the maternal-fetal interface may have downstream benefits for fetal and neonatal respiratory outcomes. Because this model induces fetal lung inflammation without fetal infection or preterm labor, it is well suited for dissecting the mechanisms by which subclinical intrauterine infection increases vulnerability to RDS and BPD, and for identifying early biomarkers and pathways that precede clinical signs of intrauterine infection or preterm labor.

## Ethics Statement

All animal procedures were approved by the Oregon Health & Science University Institutional Animal Care and Use Committee (IACUC protocol IP00000601) and conducted in accordance with the National Institutes of Health *Guide for the Care and Use of Laboratory Animals*. Animals were housed and cared for at the Oregon National Primate Research Center (ONPRC), an AAALAC International-accredited facility, under the supervision of laboratory animal veterinarians. This manuscript was prepared in accordance with the ARRIVE (Animal Research: Reporting of In Vivo Experiments) 2.0 guidelines.

## Acknowledgements

We would like to acknowledge the Oregon National Primate Research Center, Division of Comparative Medicine and specifically the Surgical Services Unit, Time-Mated Breeding, Dr Drew Martin, DVM, and Darla Jacobs for their project support. We would also like to thank the UAB Diagnostic Mycoplasma Laboratory for supplying *Ureaplasma* inoculum and performing *Ureaplasma* culture and PCR confirmation. We would like to acknowledge the Endocrine Technologies Core (ETC) and Dr. David Erickson at the Oregon National Primate Research Center (ONPRC) for providing support for multiplex and time-mated breeding hormone assays. We acknowledge the Pathology Services Unit (PSU), Integrated Pathology Core (IPC) and Clinical Pathology Laboratory at ONPRC for providing pathology support and services. Figures were created with BioRender.com.

## Supporting Information Figure Legends

S1 Fig. *UreaseB* gene detection in placenta and fetal lung. (A) Placenta (lane 5 and 6) and (B) fetal lung (lane 5) of *Ureaplasma* infected animals showed LAMP amplicons, visualized as ladder like appearance by electrophoresis on an agarose gel compared to control. Amplicon sizes are determined using 100 bp ladder run on the same agarose gel (n=4 animals/group). Control, uninfected; *Ureaplasma*, Choriodecidual *Ureaplasma parvum* infected animals.

S2 Fig. *Ureaplasma parvum* infection induced changes in inflammatory mediators in placenta. The protein level in placenta lysates are represented as mean±SEM, pg/mg protein. No changes were observed for choriodecidual *Ureaplasma* infected animals from the corresponding control (Student’s t-test/ Mann-Whitney U-test, n=3-4 animals/group).

S3 Fig. Fetal lung inflammatory marker changes. mRNA expression of *TNF-α*, *IL-6*, *MCP-1* and *VEGF* significantly upregulated and *IL-10* significantly down regulated in the fetal lung of *U.p.* animals compared to control. The mRNA expressions were normalized to the *GAPDH* housekeeping gene and represented as mean±SEM fold change. The asterisks indicate significance difference from the corresponding control (Student’s t-test/ Mann-Whitney U-test, *P<0.05, n=3-4 animals/group). Control, uninfected; *U.p.*, Choriodecidual *Ureaplasma parvum* infected animals.

S4 Fig. IL-18 associated protein changes in fetal lung. No changes were observed either at (A) mRNA expression or (B, C, D) protein levels of IL-18RAP and IL-18BP in the fetal lung of *U.p.* animals compared to control. The mRNA expression was normalized to the *GAPDH* housekeeping gene and represented as mean±SEM fold change. (B) Densitometric quantification and (C and D) representative western blot of IL-18RAP and IL-18BP in the fetal lung were normalized to GAPDH as a loading control and compared with control group. Each bar represents the mean± SEM. The asterisks indicate significance difference from the corresponding control (Student’s t-test/ Mann-Whitney U-test, n=4 animals/group). Control, uninfected; *U.p.*, Choriodecidual *Ureaplasma parvum* infected animals.

S5 Fig. MYD88 and kinase pathway changes in fetal lung. (A) mRNA expression of *MYD88*, *IRAK1*, *IRAK4*, *MAPK1* and *MAPK14* showed no change in the fetal lung of *U.p.* and control animals. The mRNA expression was normalized to the *GAPDH* housekeeping gene and represented as mean±SEM fold change. (B) Densitometric quantification and (C and D) representative western blot of MYD88, pERK1/2, Total ERK1/2, pIRAK1 (Thr209), pIRAK1 (Thr387), Total IRAK1, p p38 MAPK and Total p38 MAPK in the fetal lung showed no change in *U.p.* and control animals. Target protein levels were normalized to GAPDH as a loading control and compared with control group. The results are shown as either relative protein levels with respective to the loading control (MYD88) or as relative active phosphorylated form to total protein levels and compared with control group. Each bar represents the mean± SEM. (Student’s t-test/ Mann-Whitney U-test, n=4 animals/group). Control, uninfected; *U.p.*, Choriodecidual *Ureaplasma parvum* infected animals.

S6 Fig. Pulmonary Artery and Bronchial Airway Smooth Muscle Actin (α-SMA) expression and morphology in fetal lung. (A) Quantitative immunohistochemical analyses α-SMA staining of the pulmonary artery and bronchial airways in the fetal lung of *U.p.* animals compared to control showed no change across the groups. Each bar represents the mean± SEM. Student’s t-test/ Mann-Whitney U-test, n=3-4 animals/group. (B) Representative immunohistochemical staining for α-SMA in pulmonary artery (a, c) and bronchial airways (b, d) are shown for control (a, b) and *U.p.* (c,d) animals (Scale bar, 50 μm). Control, uninfected; *U.p.*, Choriodecidual *Ureaplasma parvum* infected animals.

S1 Table. List of loop-mediated isothermal amplication (LAMP) primers. FIP, Forward Inner Primer; BIP, Backward Inner Primer; F3, Forward Outer primer; B3, Backward Outer Primer; Loop F and Loop B, Loop Primers.

